# Validation of a Microsampling-Compatible LC-MS/MS Method for Cannabinoid Quantitation in Whole Blood

**DOI:** 10.1101/2025.09.18.676898

**Authors:** Aman A Mohammed, Mahmood Khan, Herbert Chan, Jeffrey R Brubacher

## Abstract

Recently developed dried blood sampling methods for cannabinoid quantitation require small blood volumes, making them microsampling-compatible, but have notable limitations including hematocrit-related bias for dried blood spots (DBS) and higher consumable costs for volumetric absorptive microsampling (VAMS®). To address these issues, we developed a highly sensitive liquid chromatography–tandem mass spectrometry (LC-MS/MS) method capable of quantifying cannabinoids in 50 µL of liquid whole blood, providing a practical microsampling alternative to dried blood approaches. Using liquid–liquid extraction (LLE) with sodium hydroxide alkalinization and acetonitrile precipitation, followed by quantitative analysis on an Agilent 6495 liquid chromatography-triple quadrupole (LC-TQ) mass spectrometer, we achieved lower limits of quantitation (LLOQs) of 0.10 ng/mL for Δ9-tetrahydrocannabinol (THC) and cannabinol (CBN), 0.20 ng/mL for cannabigerol (CBG), 0.30 ng/mL for cannabidiol (CBD), 0.50 ng/mL for 11-hydroxy-THC (11-OH-THC), and 5.0 ng/mL for 11-nor-9-carboxy-THC (THC-COOH). Calibration was linear from the LLOQ to 300 ng/mL for all analytes. To our knowledge, this is the first validated approach for cannabinoid quantitation in less than 100 µL of liquid whole blood with an LLOQ for THC comparable to that with the most sensitive LC-MS/MS methods using standard blood volumes. The achieved LLOQs for other cannabinoids are also suitable for forensic toxicology applications. We anticipate particular utility for obtaining evidence from suspected impaired drivers at the roadside when paired with finger-prick sampling and liquid blood microcollection tubes.This approach enables measurement of THC levels at the time of driving and thereby overcoming current limitations, including the decrease in THC levels that occurs if blood sampling is delayed, the requirement for larger sample volumes (≥100 µL), and dependence on trained phlebotomists for venipuncture.

## Introduction

Cannabis use has risen substantially in recent years, particularly following legalization in many jurisdictions, including Canada [1]. This trend has heightened concerns about cannabis-impaired driving, as Δ9-tetrahydrocannabinol (THC), the primary psychoactive compound in cannabis, is known to impair driving performance and increase collision risk [2,3]. In 2024, 16.3% of drivers involved in collisions tested positive for THC across Canada [4]. As such, many jurisdictions have established legal driving limits for THC in blood to support enforcement of impaired driving laws [5,6]. Accurate quantitation of THC in drivers is therefore critical for improving road safety, informing public policy, and enforcing traffic laws.

Despite existence of legal limits, interpreting blood THC concentrations remains challenging [7]. In current practice, blood samples from suspected impaired drivers are typically collected hours after the driving event. During this delay, THC undergoes distribution from blood into body tissue and metabolism to inactive metabolites such as 11-nor-9-carboxy-THC (THC-COOH); individual variability in cannabinoid pharmacokinetics further complicates back-extrapolation of the driver’s THC concentration at the time of driving [8]. This creates uncertainty when interpreting THC levels obtained hours after the event.

Microsampling via a fingerstick has emerged as a promising approach that could address these issues. Defined broadly as collection of ≤50 µL of blood [9], microsampling is minimally invasive, does not require trained phlebotomists, and allows for immediate roadside collection of blood, capturing cannabinoid concentrations closer to the time of driving [10]. Dried blood spot (DBS) cards and Mitra® volumetric absorptive microsampling (VAMS®) tips are among the most popular tools, offering advantages including easy storage, reduced biosafety risk, and improved analyte stability due to enzymatic inactivation during drying [11]. These features make dried microsamples highly attractive for toxicology and forensic applications.

To take advantage of microsampling techniques, there has been a growing push to develop analytical methods for dried blood sampling in toxicology [12,13]. These approaches have demonstrated sensitivity comparable to conventional liquid blood methods for cannabinoid quantitation, despite using much lower sample volumes [14-16]. This advantage comes from the use of dried matrices, such as filter paper cards or VAMS® tips, which retain cells, proteins, and lipids, significantly reducing the matrix effect that often suppresses analyte sensitivity. In addition, this feature allows analysts to bypass liquid-liquid or solid-phase extraction (SPE) steps and instead use a simple elution approach. Despite these advantages, dried blood sampling methods also have limitations. Variability in hematocrit can affect blood spreading and uniformity on cards, reducing quantitative accuracy [17]. In addition, the higher cost of VAMS® devices remains a practical barrier to widespread use [18]. As the technology is still in its early stages, no dried blood sampling methods have been implemented in forensic casework or at the roadside. For the foreseeable future, liquid blood will remain the gold standard for toxicology until these limitations are addressed, and more validation studies establish the feasibility of dried microsampling.

However, there are currently no published methods that quantitate cannabinoids from liquid blood volumes in the microsampling range (≤50 µL) with equivalent sensitivity as conventional approaches. This gap largely exists due to a combination of factors including the matrix effect associated with liquid blood, extraction-related sample loss, and the sensitivity limits of current gas and liquid chromatography-tandem mass spectrometry methods. As summarized in **Supplementary Table S1**, published liquid blood and plasma methods for cannabinoid quantitation rely on ≥100 µL of sample to achieve high sensitivity and compensate for extraction losses and matrix effects.

To address this gap, we developed a method that enables cannabinoid quantitation from only 50 µL of liquid whole blood. The method was optimized to achieve a lower limit of quantitation (LLOQ) of 0.10 ng/mL for THC, substantially lower than previously reported methods using higher sample volumes Additional analytes, including cannabinol (CBN), cannabigerol (CBG), cannabidiol (CBD), 11-hydroxy-THC (11-OH-THC), and THC-COOH, were also quantified with LLOQs suitable for forensic applications. This approach integrates microsampling with liquid blood, combining the practicality of low-volume sampling with the reliability of liquid blood analysis.

## Materials and methods

### Chemicals and Reagents

THC, CBN, CBG, CBD, 11-OH-THC, and THC-COOH analytical standards (1.0 mg/mL in methanol) and their corresponding deuterated internal standards (THC-d_3_, CBN-d_3_, CBG-d_3_, CBD-d_3_, 11-OH-THC-d_3_, THC-COOH-d_3_; 1.0 mg/mL in methanol) were obtained from Cerilliant (Round Rock, TX, USA). HPLC-grade methanol, isopropanol, acetonitrile, hexane, and ethyl acetate were purchased from VWR (Radnor, PA, USA). Liquid chromatography-mass spectrometry (LC-MS) grade formic acid and sodium hydroxide (NaOH) pellets were obtained from Fisher Scientific (Fair Lawn, NJ, USA). Deionized water was produced in-house using a Millipore Direct-Q® 5UV system (Burlington, MA, USA). Drug-free defibrinated horse blood was purchased from Dalynn (Calgary, AB, Canada) and stored at 4°C until use. Medical-grade nitrogen gas (Linde, Vancouver, BC, Canada) was used with a Reacti-Vap™ system (Fisher Scientific, Fair Lawn, NJ, USA) for solvent evaporation.

### Standards solutions preparation

A 10,000 ng/mL stock solution containing THC, CBN, CBG, CBD, 11-OH-THC, and THC-COOH was prepared in methanol and used to prepare the calibrators by serial dilution. Calibration standards were prepared by fortifying drug-free horse blood to achieve final concentrations of 0.10–300 ng/mL for THC and CBN, 0.20–300 ng/mL for CBG, 0.30–300 ng/mL for CBD, 0.50–300 ng/mL for 11-OH-THC, and 5.0–300 ng/mL for THC-COOH.

Separate methanolic stock solutions (10, 100, 1000, 10000 ng/mL) containing each cannabinoid individually were used to prepare the following blood quality controls (QCs): low (0.30 ng/mL for THC, CBN; 0.60 for CBG; 0.90 for CBD; 1.5 for 11-OH-THC; 15 for THC-COOH), mid (20 ng/mL for THC, CBN, CBG, CBD, 11-OH-THC; 120 for THC-COOH), and high (240 ng/mL for THC, CBN, CBG, CBD, 11-OH-THC, and THC-COOH).

A mixed internal standard (ISTD) solution was prepared in methanol at the following concentrations: 10 ng/mL for THC-d_3_, CBN-d_3_, CBG-d_3_, CBD-d_3_, 11-OH-THC-d_3_, and 25 ng/mL for THC-COOH-d_3_.

### Liquid-liquid extraction

50 µL of appropriate calibrator or QC was fortified with 25 µL of ISTD solution. 100 µL of 0.1 M NaOH and 100 µL of 0.1% formic acid were added sequentially and mixed for 1 minute. 500 µL of cold acetonitrile was added to precipitate proteins, followed by mixing for 5 minutes and centrifugation (4000 RPM, 10 minutes, 8^°^C). The supernatant was then transferred to clean tubes and mixed with 1.5 mL 7:3 (v/v) hexane:ethyl acetate for 10 minutes and centrifuged (4000 RPM, 10 minutes, 8^°^C). The upper organic layer was then transferred to clean tubes and evaporated at 25^°^C under nitrogen gas (flow rate of 1.5 mL/min for 5 minutes). Extracts were reconstituted in 80 µL of 75:25 (v/v) 0.1% formic acid in methanol:0.1% formic acid in water and transferred to autosampler vials.

### Instrumentation

Analyte separation was achieved using an Agilent 1290 Infinity II liquid chromatograph (Santa Clara, CA, USA) with an Agilent Poroshell 120 EC-C18 column (2.7 um, 2.1 mm x 100 mm) that was held at a temperature of 40^°^C. Mobile phase A was 0.1% formic acid in water and mobile phase B was 0.1% formic acid in methanol. The total run-time was 10.5 minutes with the following gradient profile applied: 75% B at 0.40 mL/min for 0.5 minutes, before increasing to 100% B over 4.5 minutes, hold at 100% B while increasing to 0.70 mL/min over 1 minute, hold at 0.70 mL/min for 1 min, before decreasing back to 75% B at 0.40 mL/min by 7.2 minutes and re-equilibrating for 3.3 minutes. The injection volume used was 20 µL.

Quantitative analysis was carried out on an Agilent 6495 triple quadrupole mass spectrometer. Ionization was performed by electrospray in both positive and negative polarity, and the instrument operated in dynamic multiple reaction monitoring (dMRM) mode, acquiring one quantifier and one qualifier transition per analyte and ISTD. The full list of monitored transitions is provided in **Supplementary Table S2**. The optimized MS source parameters are as follows: gas temperature (290°C); gas flow (11 L/min); nebulizer (43 psi); sheath gas temperature (370°C); sheath gas flow (12 L/min); capillary voltage (3100 V); and nozzle voltage (500 V). Agilent MassHunter Workstation (Version 12.0) was used for data acquisition and analysis.

### Method validation

Method validation followed the Scientific Working Group for Forensic Toxicology (SWGTOX) guidelines [19]. Validation criteria that were evaluated included calibration model, bias, within- and between-run precision, limit of detection, limit of quantitation, ion suppression/enhancement, interferences, carryover, dilution integrity, and processed sample stability.

The calibration model was evaluated using 7 non-zero calibrators over 5 different days. Linearity was modeled by least-squares regression with a 1/x^2^ weighting and was accepted when the residuals of individual calibrators remained within ±3 standard deviations and the correlation coefficient (R^2^) exceeded 0.99. Limits of detection (LOD) were estimated mathematically using the regression method for LOD, based on the y intercept and mean slope of the five calibration curves for each cannabinoid. The LLOQ was defined as the lowest non-zero calibrator, which was extracted in triplicate from three unique blood sources across five days (n = 15). LLOQs were accepted when qualifying ion ratios were within ±20% of the calibrators, signal-to-noise ratios were ≥10, and retention times were within ±0.1 min of the calibrators.

Carryover was investigated over 5 runs by injecting an extracted blank immediately after a 500 ng/mL fortified sample (n = 5). Carryover was considered absent if the blank response was below the signal of the lowest calibrator and no peaks were observed at the expected retention time. Bias and precision were assessed at three quality control (QC) levels across 5 days. Bias (%) was considered acceptable when within ±20% and precision (%CV) was required to be <20%. Within-run precision was calculated separately for each QC level per run, while between-run precision was determined from the mean values across all 5 days.

Ion suppression/enhancement was evaluated using post-extraction addition at the low and high QC levels. Percent matrix effect was calculated by dividing the mean peak areas of the analyte and ISTD fortified in 10 unique blood sources extracted in duplicate (n = 20) by the mean peak areas of 8 neat samples of analyte and ISTD. Acceptability was defined as <25% matrix effects and %CV <20%.

Matrix interferences were also investigated by extracting 10 unique sources of blank blood without analyte or ISTD, which allowed assessment of endogenous compounds that could contribute to background analyte signals. Responses in these blanks were considered negligible if found to be below the LLOQ and no peaks were observed at the expected retention time. Interference was further assessed by analyzing samples containing only the ISTDs (n = 3) and evaluating the signal of the analytes.

Interference was determined to be insignificant if the signal fell below the LLOQs. To examine potential ISTD crosstalk from high concentrations of cannabinoids, blank blood samples (n = 3) were fortified with 500 ng/mL of cannabinoids in the absence of ISTDs, and the deuterated transitions were monitored.

Exogenous interferences were evaluated by fortifying low QC blood samples (n = 5) with a 98-compound mixture of common impairing drugs (>5000 ng/mL) and determining if acceptable bias and precision criteria were still met (±20%). The interference mix included opioids (fentanyl, morphine, tramadol), stimulants (cocaine, MDA, methamphetamine), benzodiazepines (bromazolam, diazepam, lorazepam), antidepressants (citalopram, sertraline, venlafaxine), and other drug classes. The complete list is provided in **Supplementary Table S3**.

### Additional validation parameters

Dilution integrity was evaluated by preparing a 1:1 dilution of a blood sample fortified with 4 ng/mL of THC, CBN, CBG, CBD, and 11-OH-THC, and 100 ng/mL of THC-COOH, analyzed in triplicate over 5 days. Acceptability criteria included bias and precision within ±20%.

Processed sample stability was assessed over 72 hours in the autosampler (8^°^C). Samples fortified at the low and high QC levels were separately extracted, pooled, and aliquoted into separate autosampler vials. The first vial from each set was injected in triplicate to establish the mean baseline response (t_0_), with subsequent vials being analyzed in triplicate at 8-hour intervals. Stability was considered acceptable as long as peak areas remained within ±20% of t_0_.

## Results

### Method development

The objective of this study was to develop a highly sensitive liquid chromatography-tandem mass spectrometry (LC-MS/MS) method capable of quantifying major cannabinoids in 50 µL of whole blood, providing a feasible alternative to existing liquid and dried blood techniques.

The ionization and fragmentation parameters were optimized by individually injecting 100 ng/mL of each analyte and ISTD. Liquid-liquid extraction (LLE) was selected for sample preparation for its simplicity and cost efficiency. During method optimization, parameters such as sample pH and solvent composition were systematically adjusted to maximize the response of THC, with the response of the lowest calibrator used to gauge performance. Alkalinizing the matrix with NaOH enhanced responses for THC (+29%), CBN (+22%), CBG (+40%), CBD (+57%), and 11-OH-THC (+36%), while the THC-COOH response decreased (-6%). Incorporation of a protein precipitation step with acetonitrile further increased THC (+96%), CBN (+38%), and CBD (+61%) responses, but lowered CBG (-6%), 11-OH-THC (-46%), and THC-COOH (-76%) responses.

Different ratios of organic solvent (hexane:ethyl acetate at 9:1, 7:3, and 1:1, v/v) were compared, with the 7:3 mixture providing the most favorable and balanced partitioning for all six cannabinoids.

Reconstitution was optimized by testing several solvent ratios (100% methanol, 90:10 water:methanol 75:25 methanol:water, and 75:25 mobile phase B:mobile phase A) and evaluating the effect of mild heating (30 °C) during vortexing to improve solubility. Sequential addition of reconstitution solvent was also evaluated, where half the total volume was initially added, vortexed and transferred to the autosampler vial, before the second aliquot was added and the process was repeated. We found no increase in analyte peak of areas of the lowest calibrators when mild heat or when sequential addition was applied. The selected reconstitution condition was 80 µL of 75:25 mobile phase B:mobile phase A.

### Method validation

The calibration ranges achieved for the target cannabinoids are summarized in **Table 1**. All analytes demonstrated linear responses up to 300 ng/mL, with LLOQs corresponding to the lowest calibrators: 0.10 (THC, CBN), 0.20 (CBG), 0.30 (CBD), 0.50 (11-OH-THC), and 5.0 ng/mL (THC-COOH). Residual plots confirmed homoscedasticity, and correlation coefficients were consistently ≥0.997, supporting the use of a linear regression model. At the LLOQs, chromatographic peaks were well defined, with signal-to-noise ratios ≥10, consistent retention times, and acceptable qualifier ion ratios (±20% of calibrators). Representative chromatograms at the LLOQ are provided in **Figure 1**. Using the regression method, LODs were 0.027 (THC), 0.021 (CBN), 0.14 (CBG), 0.09 (CBD), 0.45 (11-OH-THC), and 2.2 (THC-COOH). No carryover was observed in the blank following injection of a 500 ng/mL cannabinoid sample (n = 5).

**Table 1.** Limits of detection (LOD), lower limits of quantitation (LLOQ), and calibration results for cannabinoids.

| Analyte | Internal Standard | LOD (ng/mL) | LLOQ (ng/mL) | Slope ± SD (n = 5) | y-int ± SD (n = 5) | R<sup>2</sup> ± SD (n = 5) | Linear Range (ng/mL) |
| --- | --- | --- | --- | --- | --- | --- | --- |
| THC | THC-d<sub>3</sub> | 0.027 | 0.10 | 0.044 ± 0.002 | -0.001 ± 0.000 | 1.000 ± 0.000 | 0.10–300 |
| CBN | CBN-d<sub>3</sub> | 0.031 | 0.10 | 0.291 ± 0.004 | 0.008 ± 0.003 | 0.999 ± 0.000 | 0.10–300 |
| CBG | CBG-d<sub>3</sub> | 0.14 | 0.20 | 0.127 ± 0.005 | 0.002 ± 0.006 | 0.999 ± 0.000 | 0.20–300 |
| CBD | CBD-d<sub>3</sub> | 0.09 | 0.30 | 0.134 ± 0.007 | 0.002 ± 0.003 | 0.999 ± 0.000 | 0.30–300 |
| 11-OH-THC | 11-OH-THC-d<sub>3</sub> | 0.45 | 0.50 | 0.213 ± 0.061 | 0.029 ± 0.030 | 0.999 ± 0.000 | 0.50–300 |
| THC-COOH | THC-COOH-d<sub>3</sub> | 2.2 | 5.0 | 0.0418 ± 0.084 | 0.008 ± 0.028 | 0.997 ± 0.001 | 5.0–300 |

**Figure 1.**
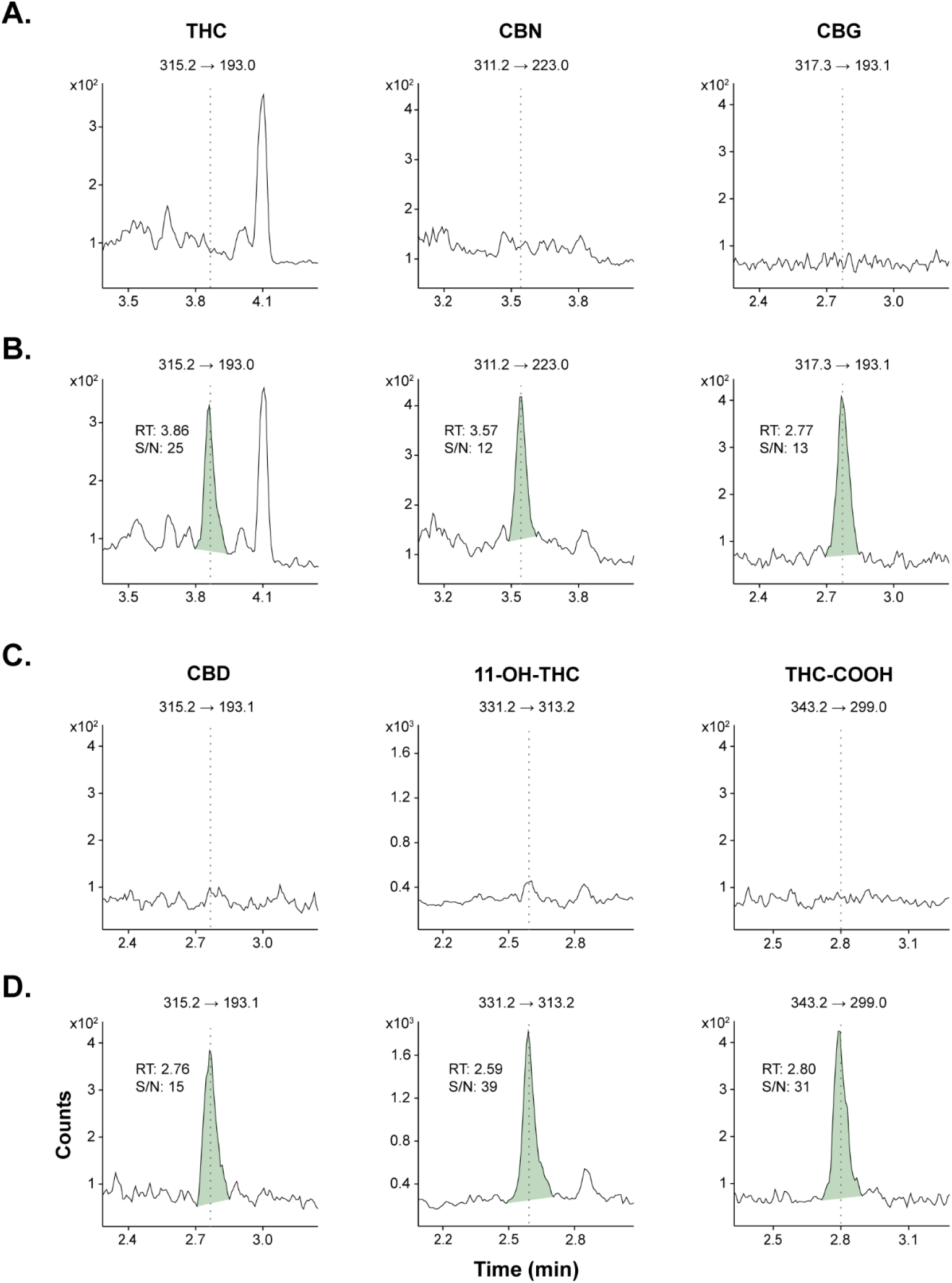
Ion chromatograms from an extracted blank blood matrix for (**A**) Δ9-tetrahydrocannabinol (THC), cannabinol (CBN), cannabigerol (CBG) and (**C**) cannabidiol (CBD), 11-hydroxy-THC (11-OH-THC), 11-nor-9-carboxy-THC (THC-COOH). Ion chromatograms from an extracted standard at the limit of quantitation for (**B**) THC (0.10 ng/mL), CBN (0.10 ng/mL), CBG (0.20 ng/mL) and (**D**) CBD (0.30 ng/mL), 11-OH-THC (0.50 ng/mL), and THC-COOH (5.0 ng/mL)

Bias and precision were evaluated by fortifying drug-free horse blood with each cannabinoid at low, mid, and high concentrations over 5 days (n = 25). The low QC level for each cannabinoid was established as 3 times the LLOQ, with the high QC level being 80% of upper limit of quantitation (ULOQ). The mid QC level was set at 120 ng/mL for THC-COOH and 20 ng/mL for the other cannabinoids to better reflect expected concentrations, as THC-COOH levels tend to remain elevated for extended periods compared to other cannabinoids [20,21]. Across all cannabinoids and QC concentrations, bias ranged from -14.0% to 16.1%. Within-run precision was between 0.9% to 14.5%, and between-run precision ranged from 1.4% to 8.6%. All data ranges met the 20% acceptance criteria of the SWGTOX guidelines. Dilution integrity was also assessed with a 2-fold dilution of a blank fortified with 4 ng/mL of THC, CBN, CBG, CBD, and 11-OH-THC, and 100 ng/mL of THC-COOH (n = 25). Acceptable bias and precision criteria were met. **Table 2** details the bias and precision results for each cannabinoid for each QC concentration, the dilution integrity study, and the LLOQs.

**Table 2.**
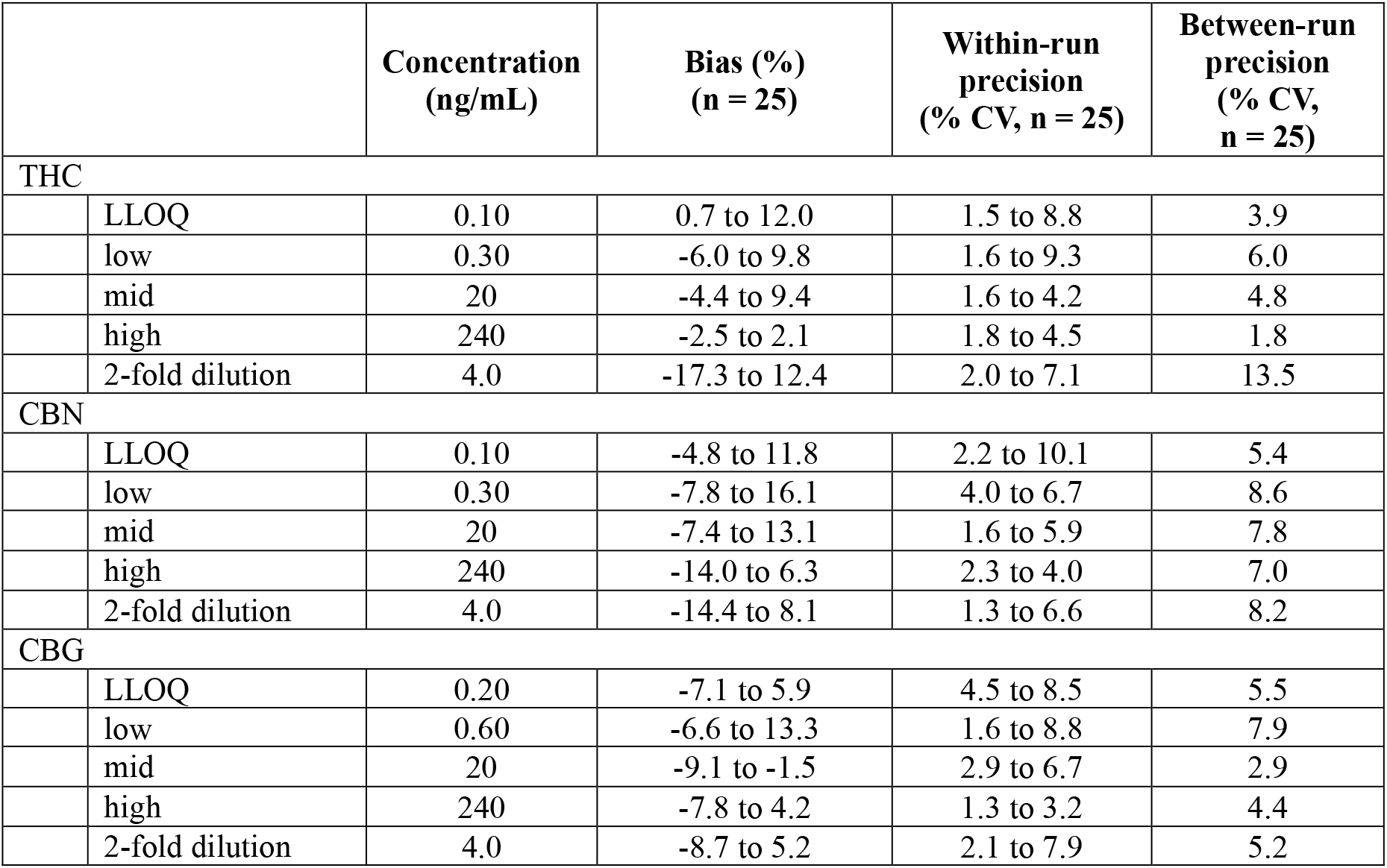
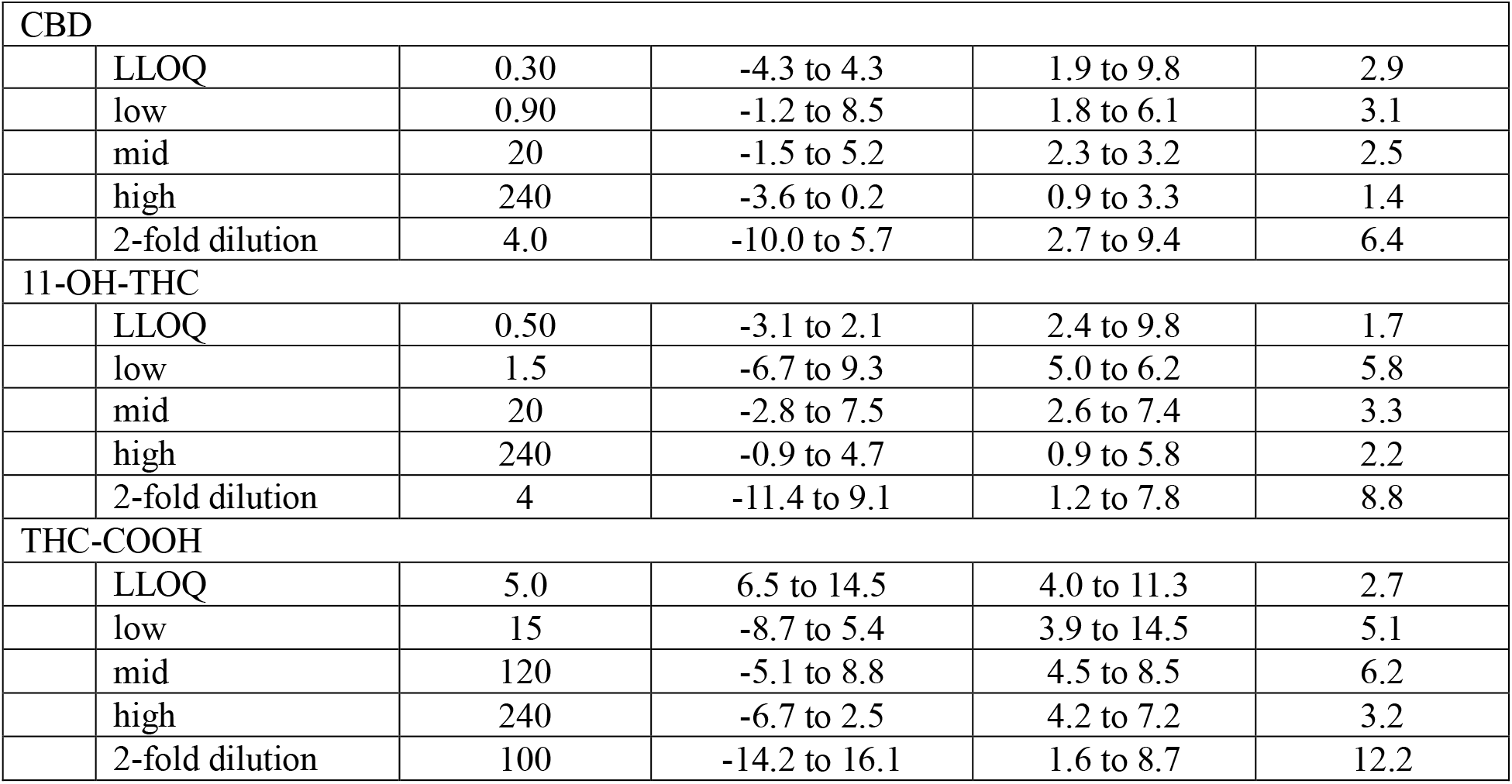
Bias and precision data of lower limits of quantitation (LLOQ), quality controls (low, mid, high) and 2-fold dilution samples.

|  | Concentration (ng/mL) | Bias (%) (n = 25) | Within-run precision (% CV, n = 25) | Between-run precision (% CV, n = 25) |
| --- | --- | --- | --- | --- |
| THC |  |  |  |  |
| LLOQ | 0.10 | 0.7 to 12.0 | 1.5 to 8.8 | 3.9 |
| low | 0.30 | -6.0 to 9.8 | 1.6 to 9.3 | 6.0 |
| mid | 20 | -4.4 to 9.4 | 1.6 to 4.2 | 4.8 |
| high | 240 | -2.5 to 2.1 | 1.8 to 4.5 | 1.8 |
| 2-fold dilution | 4.0 | -17.3 to 12.4 | 2.0 to 7.1 | 13.5 |
| CBN |  |  |  |  |
| LLOQ | 0.10 | -4.8 to 11.8 | 2.2 to 10.1 | 5.4 |
| low | 0.30 | -7.8 to 16.1 | 4.0 to 6.7 | 8.6 |
| mid | 20 | -7.4 to 13.1 | 1.6 to 5.9 | 7.8 |
| high | 240 | -14.0 to 6.3 | 2.3 to 4.0 | 7.0 |
| 2-fold dilution | 4.0 | -14.4 to 8.1 | 1.3 to 6.6 | 8.2 |
| CBG |  |  |  |  |
| LLOQ | 0.20 | -7.1 to 5.9 | 4.5 to 8.5 | 5.5 |
| low | 0.60 | -6.6 to 13.3 | 1.6 to 8.8 | 7.9 |
| mid | 20 | -9.1 to -1.5 | 2.9 to 6.7 | 2.9 |
| high | 240 | -7.8 to 4.2 | 1.3 to 3.2 | 4.4 |
| 2-fold dilution | 4.0 | -8.7 to 5.2 | 2.1 to 7.9 | 5.2 |

|  |  |  |  |  |  |
| --- | --- | --- | --- | --- | --- |
| CBD |  |  |  |  |  |
|  | LLOQ | 0.30 | -4.3 to 4.3 | 1.9 to 9.8 | 2.9 |
|  | low | 0.90 | -1.2 to 8.5 | 1.8 to 6.1 | 3.1 |
|  | mid | 20 | -1.5 to 5.2 | 2.3 to 3.2 | 2.5 |
|  | high | 240 | -3.6 to 0.2 | 0.9 to 3.3 | 1.4 |
|  | 2-fold dilution | 4.0 | -10.0 to 5.7 | 2.7 to 9.4 | 6.4 |
| 11-OH-THC |  |  |  |  |  |
|  | LLOQ | 0.50 | -3.1 to 2.1 | 2.4 to 9.8 | 1.7 |
|  | low | 1.5 | -6.7 to 9.3 | 5.0 to 6.2 | 5.8 |
|  | mid | 20 | -2.8 to 7.5 | 2.6 to 7.4 | 3.3 |
|  | high | 240 | -0.9 to 4.7 | 0.9 to 5.8 | 2.2 |
|  | 2-fold dilution | 4 | -11.4 to 9.1 | 1.2 to 7.8 | 8.8 |
| THC-COOH |  |  |  |  |  |
|  | LLOQ | 5.0 | 6.5 to 14.5 | 4.0 to 11.3 | 2.7 |
|  | low | 15 | -8.7 to 5.4 | 3.9 to 14.5 | 5.1 |
|  | mid | 120 | -5.1 to 8.8 | 4.5 to 8.5 | 6.2 |
|  | high | 240 | -6.7 to 2.5 | 4.2 to 7.2 | 3.2 |
|  | 2-fold dilution | 100 | -14.2 to 16.1 | 1.6 to 8.7 | 12.2 |

Matrix effects were examined by post-extraction addition at the low and high QC levels. Ion suppression/enhancement across 10 unique blood matrices was ≤±25% with %CV <20%, indicating effective removal of interferents during extraction.

Extracted blank blood samples from 10 unique blood sources (n = 20) were found to be free of interferents. Endogenous interferences were qualitatively evaluated by monitoring quantifier transitions following injection of extracted blank blood samples fortified with only ISTDs (n = 3). Additionally, blank blood samples (n = 3) fortified with cannabinoids at the ULOQ but without ISTD were assessed. Across these samples, no crosstalk between deuterated and non-deuterated cannabinoids was found.

Extracted blood samples containing cannabinoids at the low QC level (n = 5) were fortified with 98 common impairing drugs to assess exogenous interferences. Accuracy and precision were within 20% and thus concluded to be acceptable.

Stability was evaluated for extracted samples at low and high QC levels stored in the autosampler (8^°^C) for 72 hours. Results expressed as the percent difference from t_0_ peak areas are shown in **Supplementary Table S4**. All analytes remained stable (±20% of t_0_ peak area) after 24 hours. At 48 hours, CBN, CBG,

11-OH-THC, and THC-COOH fell below 80% of the t_0_ peak area at one QC level. By 72 hours, THC, CBN, 11-OH-THC, and THC-COOH were within acceptance criteria, while CBG and CBD at the low QC level showed >20% loss relative to t_0_.

## Discussion

To our knowledge, this study presents the first validated method capable of quantifying THC at 0.10 ng/mL using 50 µL of whole blood, while simultaneously quantifying CBN, CBG, CBD, 11-OH-THC, and THC-COOH. Previously published methods have required ≥100 µL of blood [22-24], relied on the cleaner matrix of plasma to increase sensitivity [25-27], employed SPE [28-30], or a combination of these strategies to reach sub-ng/mL sensitivity. Some approaches also use separate acidic and basic extractions to optimize recovery of both THC and THC-COOH [31-33]. Our method incorporates both NaOH and formic acid in a single extraction, providing a flexible point of adjustment. Increasing basicity enhances extraction efficiency for THC, which is particularly relevant when testing for impairment. Conversely, more acidic conditions can improve recovery of THC-COOH, which can be more informative for assessing cannabis use history. In this study, the method was optimized for THC, reflecting its envisioned application in roadside microsampling, where timely collection provides more accurate estimates of THC levels at the time of collision or other driving incident.

While the use of SPE is well established for improving sensitivity through reduced analyte loss and enhanced matrix clean-up [34], our LLE method achieves superior LLOQs for THC without the need for SPE. This is particularly advantageous for high throughput applications, as the high cost of solid phase cartridges can be a practical barrier. Although our LLOQs for THC metabolites THC-COOH and 11-OH-THC are higher than those reported in some methods [24,27], they are consistent with several recent studies [22,23,29] and remain suitable for quantitation. Including 11-OH-THC is particularly important, as it is psychoactive and often reaches higher blood concentrations after oral cannabis consumption compared to inhalation [35]. Further, our method also validates the quantitation of additional cannabinoids, CBN, CBG, and CBD, which are often excluded from published methods. Although these compounds are non-psychoactive and less studied, they may provide valuable contextual information.

CBG, known as the “mother cannabinoid” for its role as the precursor of other cannabinoids in the cannabis plant, undergoes rapid hepatic metabolism [36] and may therefore indicate recent consumption. CBN, which forms from the oxidative degradation of THC in the cannabis plant, could provide insight into the type or age of product consumed [37]. CBD measurement is also increasingly relevant, given the widespread availability of CBD-rich cannabis products and edibles, as well as its growing use for medical purposes [38,39]. Together, inclusion of these cannabinoids extends interpretive power beyond THC alone and may strengthen forensic assessments of cannabis use.

The present method not only offers a robust alternative to existing liquid blood assays but also provides a practical counterpart to dried blood sampling approaches. By using only 50 µL of whole blood, it preserves the advantages of microsampling while avoiding key limitations associated with DBS and VAMS®, such as hematocrit-related variability and higher consumable costs. In our experience, obtaining 50 µL of blood via a fingerstick is feasible with appropriate lancets, and collection into microtubes such as Sarstedt Microvette® devices can provide a simple, low-cost option for storage. Moreover, we demonstrate that a 1:1 dilution can be performed without compromising accuracy, supporting the feasibility of using as little as 25 µL of liquid blood when smaller volumes are collected.

Another important distinction lies in sample preparation. DBS and VAMS® methods rely on protein precipitation with methanol or acetonitrile [14-16], taking advantage of the partial retention of cells and proteins within dried matrices to reduce matrix complexity and achieve low LLOQs. In contrast, liquid blood methods generally incorporate an additional organic separation step, which helps remove endogenous material. However, phospholipids, particularly lysophosphatidylcholines and lysophosphatidylethanolamines, can co-elute with THC in LC-MS workflows and have been shown to cause pronounced ion suppression of cannabinoids in acetonitrile extracts of whole blood [40,41]. The long-term impact of residual phospholipid co-extraction from DBS and VAMS® extracts on instrument performance has not yet been characterized. Future studies of dried blood sampling should therefore include assessments of instrument robustness, including evaluations of sensitivity and column backpressure over extended injection series (>1000 injections) in comparison with conventional organic extraction workflows.

A major advantage of dried blood sampling is the ability to store samples at room temperature, since water loss during drying halts most enzymatic degradation reactions. Recent studies have shown that THC, 11-OH-THC, and THC-COOH remain stable on DBS cards for up to three months at room temperature [14], while VAMS® devices maintain stability for at least 30 days [15]. For liquid blood (≥2 mL), however, the available stability data are much older, with reports from the 1980s indicating that cannabinoids remain stable for approximately one month in glass tubes at room temperature [42], whereas storage in plastic containers leads to markedly greater analyte loss [43]. Given advances in collection devices designed for very low blood volumes [44], updated stability studies are needed to confirm storage conditions suitable for microsampling applications.

Practical limitations to consider when utilizing our method include the challenge of working with low blood volumes. When only °50 µL of blood is available, accurately aliquoting the entire amount be difficult during forward pipetting using positive-displacement pipettes, as bubbles may form. However, our results demonstrate that sample dilution is possible without compromising accuracy or precision. It should also be emphasized that although our extraction procedure has been highly optimized, the results reported here were achieved using a state-of-the-art Agilent 6495 liquid chromatography-triple quadrupole mass spectrometer, which has an instrument detection limit of 0.30 fg of reserpine when injected onto the column [45]. Comparable results may therefore depend on access to instrumentation of similar sensitivity.

## Conclusion

The present LC-MS/MS method enables quantitation of six major cannabinoids of forensic interest: THC, CBN, CBG, CBD, 11-OH-THC, and THC-COOH. Through the use of only 50 µL of liquid whole blood and an LLE procedure with LLOQs comparable to the most sensitive published studies, this method represents an advance in liquid blood analysis of cannabinoids and provides a feasible alternative to dried blood sampling approaches that could be used on the roadside for measurement of THC levels at the time of driving.

## Funding

Funding for this research was provided by the Canadian Institutes of Health Research [375264].

## Supplementary data

The online version contains supplementary data.

## Notes

### Competing Interest Statement

The authors have declared no competing interest.

