## Supplementary Data for "Validation of a Microsampling-Compatible LC-MS/MS Method for Cannabinoid Quantitation in Whole Blood"

**Supplementary Table S1**. Published methods for cannabinoid quantitation in whole blood, plasma, or serum. The table summarizes sample volume, extraction procedure, analytical technology, and reported lower limits of quantitation (LLOQ) for Δ9-tetrahydrocannabinol (THC), 11-nor-9-carboxy-THC (THC-COOH), 11-hydroxy-THC (11-OH-THC), cannabinol (CBN), cannabigerol (CBG), and cannabidiol (CBD). References for each method are listed at the end of the Supplementary Data.

| Reference | Sample (µL) | Extraction Procedure | Analytical Technology | LLOQ (ng/mL) | | | | | |
|---|---|---|---|---|---|---|---|---|---|
| | | | | THC | THC-COOH | 11-OH-THC | CBN | CBG | CBD |
| Proença et al., 2023 [1] | 100 WB | Protein precipitation | LC-MS/MS | 0.5 | 0.5 | 0.5 | - | - | - |
| Hädener et al., 2016 [2] | 100 WB | Protein precipitation | LC-MS/MS | - | 5 | - | - | - | - |
| Hubbard et al., 2020 [3] | 200 WB | SPE | LC-MS/MS | 0.5 | 1 | 1 | 0.5 | 1 | 0.5 |
| Scheidweiler et al., 2016 [4] | 200 WB | SPE | LC-MS/MS | 0.5 | 0.5 | 0.5 | 0.5 | 1 | 0.5 |
| Orfanidis et al., 2020 [5] | 200 WB | QuEChERS (dSPE) | LC-MS/MS | 18.54 | 3.31 | - | - | - | - |
| Palazzoli et al., 2018 [6] | 200 WB | Protein precipitation | LC-MS/MS | 1 | 1 | 1 | - | - | 0.5 |
| Fernandez et al., 2008 [7] | 250 WB | LLE | LC-MS/MS | 0.5 | 2 | 1 | - | - | - |
| Chan-Hosokawa et al., 2022 [8] | 250 WB | LLE | LC-MS/MS | 0.5 | 5 | 1 | - | - | - |
| Frei et al., 2022 [9] | 250 WB | SPE | GC-MS/MS | 0.3 | 3 | 0.3 | 0.2 | - | 0.3 |
| Thomas et al., 2007 [10] | 500 WB | LLE | GC-MS/MS | 0.5 | 2.5 | 0.5 | - | - | - |

**Supplementary Table S1 Continued**

| Reference | Sample (µL) | Extraction Procedure | Analytical Technology | LLOQ (ng/mL) | | | | | |
|---|---|---|---|---|---|---|---|---|---|
| | | | | THC | THC-COOH | 11-OH-THC | CBN | CBG | CBD |
| Simoes et al., 2011 [11] | 500 WB | SPE | LC-MS/MS | 0.5 | 0.5 | 0.5 | - | - | - |
| Schwope et al., 2011 [12] | 500 WB | SPE | LC-MS/MS | 1 | 1 | 1 | 1 | - | 1 |
| Holland et al., 2011 [13] | 500 WB | SPE | GC-MS | 0.5 | 0.5 | 0.5 | - | - | - |
| Castro et al., 2018 [14] | 500 WB | SPE | GC-MS/MS | 0.93 | 0.99 | 0.87 | - | - | - |
| Ferrari et al., 2022 [15] | 500 WB | QuEChERS (dSPE) | LC-MS/MS | 4 | 10 | - | - | - | - |
| Joye et al., 2020 [16] | 1000 WB | LLE | LC-MS/MS | 0.4 | 2.5 | 0.4 | - | - | 0.4 |
| Andrews et al., 2012 [17] | 1000 WB | LLE | GC-MS | 0.25 | 0.5 | 0.25 | 0.25 | - | 0.5 |
| Reber et al., 2022 [18] | 1000 WB | SPE | LC-MS/MS | 1 | 5 | - | - | - | - |
| Teixeira et al., 2007 [19] | 1000 WB | SPE | LC-MS | 2 | 2 | 25 | - | - | - |
| Giroud et al., 2001 [20] | 1000 WB | SPE | GC-MS/MS | 1 | 5 | 1 | - | - | - |
| Hoffman et al., 2020 [21] | 2000 WB | SPE | GC-MS | 1 | 1 | - | - | - | - |
| da Silva et al., 2020 [22] | 100 P | LLE | LC-MS/MS | 0.5 | 1 | 0.5 | 0.5 | - | 0.5 |
| Manca et al., 2022 [23] | 100 P | Protein precipitation | LC-MS/MS | 5 | 5 | - | - | - | - |

**Supplementary Table S1 Continued**

| Reference | Sample (µL) | Extraction Procedure | Analytical Technology | LLOQ (ng/mL) | | | | | |
|---|---|---|---|---|---|---|---|---|---|
| | | | | THC | THC-COOH | 11-OH-THC | CBN | CBG | CBD |
| Sempio et al., 2021 [24] | 200 P | Protein precipitation | LC-MS/MS | 0.78 | 0.78 | 3.13 | 0.78 | 0.78 | 0.78 |
| Rosado et al. 2017 [25] | 250 P | MEPS (SPE) | GC-MS/MS | 0.1 | 0.1 | 0.1 | - | - | - |
| Andrenyak et al., 2017 [26] | 1000 P | LLE | GC-MS/MS | 0.1 | 0.5 | 0.1 | - | - | 0.25 |
| Álvarez-Freire et al., 2023 [27] | 1000 P | LLE | GC-MS/MS | 50 | 80 | - | - | - | - |
| Pichini et al., 2020 [28] | 100 S | LLE | LC-MS/MS | 0.12 | 0.19 | 0.12 | - | - | 0.13 |

WB – whole blood; P – plasma; S – serum; LLE – liquid-liquid extraction; SPE – solid phase extraction; dSPE – dispersive SPE; LC-MS/MS – liquid chromatography-tandem mass spectrometry; GC-MS/MS – gas chromatography-tandem mass spectrometry; MEPS - microextraction by packed sorbent.

**Supplementary Table S2.** Optimized mass transitions of analytes and internal standards.

| Compound | RT | Type | Precursor Ion | Product Ion | CE | Polarity |
|---|---|---|---|---|---|---|
| THC | 3.86 | Quantifier | 315.2 | 193.0 | 24 | Positive |
| | | Qualifier | 315.2 | 123.0 | 37 | Positive |
| THC-$d_3$ | 3.85 | Quantifier | 318.3 | 196.1 | 30 | Positive |
| | | Qualifier | 318.3 | 123.0 | 40 | Positive |
| CBN | 3.57 | Quantifier | 311.2 | 223.0 | 24 | Positive |
| | | Qualifier | 311.2 | 293.2 | 16 | Positive |
| CBN-$d_3$ | 3.56 | Quantifier | 314.2 | 296.1 | 20 | Positive |
| | | Qualifier | 314.2 | 223.1 | 20 | Positive |
| CBG | 2.78 | Quantifier | 317.3 | 193.1 | 20 | Positive |
| | | Qualifier | 317.3 | 123.0 | 20 | Positive |
| CBG-$d_3$ | 2.77 | Quantifier | 320.3 | 196.0 | 14 | Positive |
| | | Qualifier | 320.3 | 123.0 | 36 | Positive |
| CBD | 2.77 | Quantifier | 315.2 | 193.1 | 34 | Positive |
| | | Qualifier | 315.2 | 123.0 | 20 | Positive |
| CBD-$d_3$ | 2.76 | Quantifier | 318.2 | 196.1 | 20 | Positive |
| | | Qualifier | 318.2 | 123.2 | 30 | Positive |
| 11-OH-THC | 2.56 | Quantifier | 331.2 | 313.2 | 14 | Positive |
| | | Qualifier | 331.2 | 193.1 | 5 | Positive |
| 11-OH-THC-$d_3$ | 2.56 | Quantifier | 334.2 | 316.2 | 10 | Positive |
| | | Qualifier | 334.2 | 196.1 | 20 | Positive |
| THC-COOH | 2.81 | Quantifier | 343.2 | 299.0 | 20 | Negative |
| | | Qualifier | 345.2 | 299.1 | 20 | Negative |
| THC-COOH-$d_3$ | 2.80 | Quantifier | 348.2 | 302.2 | 20 | Positive |
| | | Qualifier | 348.2 | 330.2 | 10 | Positive |

**Supplementary Table S3.** List of 98 common impairing drugs analyzed for exogenous interference study.

| Category | Compounds Tested |
|---|---|
| Alcohol | ethanol |
| Opioids | 6-acetylmorphine, buprenorphine, carfentanil, codeine, EDDP, etodesnitazene, fentanyl, hydrocodone, hydromorphone, isotonitazene, meperidine, methadone, mitragynine, morphine, norfentanyl, ortho-methylfentanyl, ortho-fluorofentanyl, oxcodone, para-fluorofentanyl, protonitazene, tramadol |
| CNS Stimulants | amphetamine, benzoylecgonine, cocaethylene, cocaine, MDA, MDMA, methamphetamine |
| Dissociative Anesthetics | dextromethorphan, ketamine, phencyclidine |
| Benzodiazepines | 7-aminoclonazepam, 7-aminoflunitrazepam, 7-aminonitrazepam, 8-aminoclonazolam, alprazolam, bromazolam, chlordiazepoxide, chlorpheniramine, clonazepam, desalkylflurazepam, desalkylgidazepam, deschloroetizolam, diazepam, etizolam, flualprazolam, flubromazepam, flunitrazepam, lorazepam, nitrazepam, nordiazepam, oxazepam, temazepam, zolpidem, zopiclone |
| Antidepressants | amitriptyline, bupropion, citalopram, clomipramine, desipramine, doxepin, fluoxetine, hydroxybupropion, imipramine, mirtazapine, didesmethylcitalopram, norsertraline, nortriptyline, o-desmethylvenlafaxine, paroxetine, sertraline, trazodone, venlafaxine |
| Anticonvulsants | carbamazepine, gabapentin, lamotrigine, levetiracetam, phenytoin, topiramate |
| Antipsychotics | aripiprazole, chlorpromazine, clozapine, haloperidol, loxapine, olanzapine, quetiapine, risperidone, ziprasidone, |
| Antihistamines | cetirizine, cyclobenzaprine, diphenhydramine, doxylamine, |
| Hallucinogen | LSD, psilocin |
| Other | medetomidine, naloxone, xylazine |

**Supplementary Table S4.** Stability of extracted cannabinoid samples at the low and high quality control levels following storage in the autosampler at 8ºC for 24, 48, and 72 hours.

| | **24 h autosampler (% difference, n = 3)** | | **48 h autosampler (% difference, n = 3)** | | **72 h autosampler (% difference, n = 3)** | |
|---|---|---|---|---|---|---|
| **Analyte** | **Low** | **High** | **Low** | **High** | **Low** | **High** |
| THC | -15.4 | -12.0 | -19.7 | -17.8 | -13.2 | -16.9 |
| CBN | -17.2 | -16.4 | -18.9 | -24.0 | -17.4 | -19.5 |
| CBG | -11.7 | -19.8 | -19.9 | -26.5 | -35.6 | -18.1 |
| CBD | -0.4 | -8.2 | -14.0 | -19.7 | -27.3 | -8.4 |
| 11-OH-THC | -13.6 | -11.8 | -17.1 | -21.8 | -18.4 | -15.8 |
| THC-COOH | -17.7 | -1.9 | -24.1 | -16.2 | -15.4 | -12.1 |
